# A single-cell atlas of intestinal immune cells across the day-night cycle reveals dynamic populations

**DOI:** 10.64898/2026.01.24.701519

**Authors:** Robert W. Maples, Gabriella Quinn, Tarun Srinivasan, Chaitanya Dende, Lora V. Hooper, Julie K. Pfeiffer

## Abstract

The small intestine houses an array of immune cells that receive diverse inputs from food intake, microbiota, and other cues that vary by time of day. However, how diurnal variation influences intestinal immune cell proportions and functions is unclear. Here, we use flow cytometry and single cell RNA sequencing to establish an atlas of 815,073 mouse small intestine immune cells at four times across the day-night cycle. These data suggest possible temporal coordination of dendritic cell antigen processing and subsequent T cell antigen recognition. Most cells express circadian clock genes and have intrinsic oscillatory transcriptomes. However, differentiated antibody-producing plasma cells have minimal circadian gene expression and instead may receive extrinsic oscillatory cues from other cell types. Finally, certain populations of B cells are extremely dynamic, with broad transcriptional changes within a six hour time span. This dataset provides insight into the circadian dynamics of intestinal immunity.

**Summary:**

- An atlas of 815,073 small intestine immune cells across four time-points reveals a large proportion of naïve B and T cells.
- Gene expression profiles suggest coordination of antigen processing in dendritic cells prior to antigen recognition by T cells.
- Th17 and innate lymphoid cells have high expression of circadian clock genes and most immune cells have rhythmic gene expression.
- Populations of certain B cell subtypes, including transitional B cells and centrocytes, are extremely dynamic with large shifts over a six hour time frame.
- Terminally differentiated antibody-producing plasma cells have minimal circadian gene expression and few oscillatory genes.

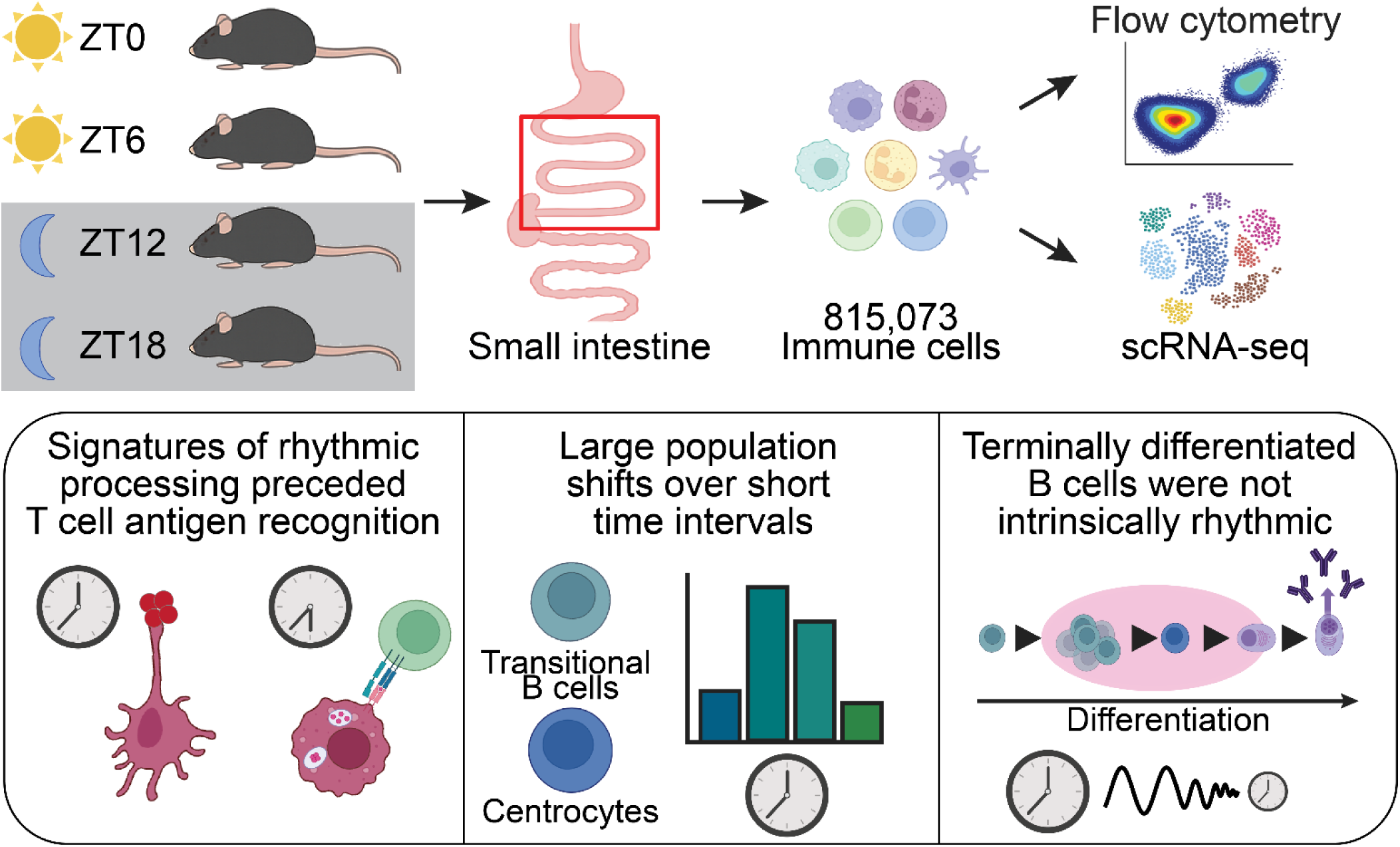

## Introduction

Mice have variable susceptibility to pathogens depending on time of day, and temporal regulation of host immune responses contributes to differential infection outcomes^1–3^. Over sixty years ago, it was recognized that susceptibility of mice to lipopolysaccharide treatment varied by time of day^4^. Since then, mechanisms underlying diurnal immune variation have been explored including circadian control of innate responses in macrophages^5^, immune cell trafficking to lymph nodes and tissues^6–8^, and differentiation pathways in Th17 cells and type 3 innate lymphoid cells (ILC3)^9–12^. More recent studies have revealed diurnal regulation of mucosal and intestinal biology. For example, microbiota aid diurnal coordination of innate immunity through a myeloid-ILC3 pathway^13^ and type 2 innate lymphoid cells (ILC2) respond to feeding cues in anticipation of potential food-borne pathogens^14^. Additionally, IgA protein levels are rhythmic in the mouse intestine^15^. However, there is limited information on whether populations of immune cells oscillate in the intestine and whether oscillatory gene expression is broad or occurs in just a subset of immune cells.

The gastrointestinal tract has many immune cells and is an important site for their maturation and differentiation in response to diverse microbial and dietary cues.

Intestinal immune cells originate from fetal or bone marrow derived sources^16–19^ and reside in the lamina propria, which underlies a single layer of epithelial cells. Within the lamina propria, many immune cells are located within gut-associated lymphoid tissue (GALT), including Peyer’s patches and isolated lymphoid follicles. B cell differentiation occurs in germinal centers of Peyer’s patches with dendritic cell (DC) and T cell help.

In mammals, diurnal cycles generate circadian rhythms that are organized in peripheral tissues through endocrine and neural pathways emanating from the brain^20–23^. The core oscillator is comprised of at least three interlocking feedback loops, including transcription factors CLOCK and BMAL1 that drive expression of their own repressors, PER and CRY proteins^24,25^. Many important studies on circadian biology have focused on the brain and liver, revealing rhythmic control of physiology and metabolism, as well as rhythmic genes^26,27^. The immune system is also regulated by circadian rhythms, including immune cell development and innate and adaptive responses^5–8,28^. However, the influence of diurnal cycles on broad immune cell populations is unclear.

To better understand the scale and diversity of immune cells in the small intestine, we examined their populations at four time points throughout the day-night cycle in mice. Initially, we used flow cytometry with a 12-marker panel. While we observed rhythmicity in some cell populations, the limited number of markers was insufficient to capture the full diversity of immune cells in the small intestine. Therefore, we performed single-cell RNA sequencing (scRNA-seq) to profile over 800,000 cells across time. We identified 33 distinct cell types and examined their gene expression patterns over time. We found time-dependent expression of DC antigen processing genes prior to transcription of T cell antigen recognition gene signatures. Additionally, we observed differential circadian gene expression among cell types, with high expression and oscillation in certain T cell and innate lymphoid cell populations but low expression and oscillation in terminally differentiated B cell lineages including antibody-producing plasmablasts and plasma cells. Finally, two populations of B cells—centrocytes and transitional B cells—were very dynamic, with large population shifts over a six hour window. Overall, these data create the largest small intestine immune cell atlas to date, provide an extensive list of marker genes for 33 types of immune cells, reveal oscillatory versus non-oscillatory genes, and illuminate which cell types are under circadian gene regulation in the small intestine.

## Methods

### Mice

For flow cytometry experiments, C57BL/6J (Strain# 000664) mice were from Jackson Labs or colony-maintained mice. For scRNA-seq experiments, only Jackson Labs C57BL/6J mice were used. All experiments used male mice 9–11-weeks old that were housed in 12-hour light/ 12-hour dark cycle in a specific pathogen-free barrier facility prior to experiments. Within a conventional barrier animal housing facility, modulations of lighting conditions occurred in isolation cabinets with green LED lights programmed by Clock Labs v3.604 and Chamber Control Software v4.114 (Actimetrics Inc., Wilmette, IL). Mice were acclimatized for 2 weeks for flow cytometry experiments or 3 weeks for scRNA-seq experiments.

### Lamina propria cell preparations

Small intestines were harvested at designated ZT and lamina propria immune cells were collected following a modified previously published method^11,12^. Briefly, fat-stripped intestines (including Peyer’s patches) were treated chemically and enzymatically with mechanical agitation to separate the epithelial layer and the extracellular matrix to isolate lamina propria-derived immune cells. However, modifications were made to further enhance the retrieval of immune cells. Peyer’s patches were not removed and two sequential 15-minute incubations with Hank’s buffered salt solution (HBSS, Thermo Fisher, 14170112) containing 10 mM EDTA, 5% FBS, and 4.71 mM sodium bicarbonate were used to strip the epithelial layer. Furthermore, after enzymatic treatments, cells were filtered using 70µm conical strainers. Finally, a 40% Percoll gradient was used to separate immune cells, which were in the pellet after centrifugation.

### Flow Cytometry

Isolated immune cells from lamina propria preparations were initially washed and resuspended in PBS into a 96 well V-bottom an untreated tissue culture plate. Cells were then stained with 1:1000 Zombie Aqua viability stain following manufacturer’s protocol. Cells were pelleted at 400 g for 5 min and then resuspended in a 1:1000 αCD16/32 cocktail for 15 min to prevent Fc-receptor binding. Cells were washed with FACS buffer (1x PBS and 3% FBS) and resuspended in a combined solution of antibodies listed in **Table 1** at the appropriate dilutions for 10 min protected from light. After washing, cells were treated with FACs buffer and stained with antibodies against cell surface markers for distinction of cells listed in **Table 1**. Cells were then fixed using 4% paraformaldehyde for 30 min at room temperature protected from light, washed in FACs buffer, and immediately analyzed using a Novocyte 3005. Antibody binding compensation beads (Fisher Scientific, 50-112-9040) and amine reactive beads (Thermo Fisher, A10346) were used to establish voltages and compensation for multicolor flow cytometry. The data were evaluated with NovoExpress software.

**Table 1.**
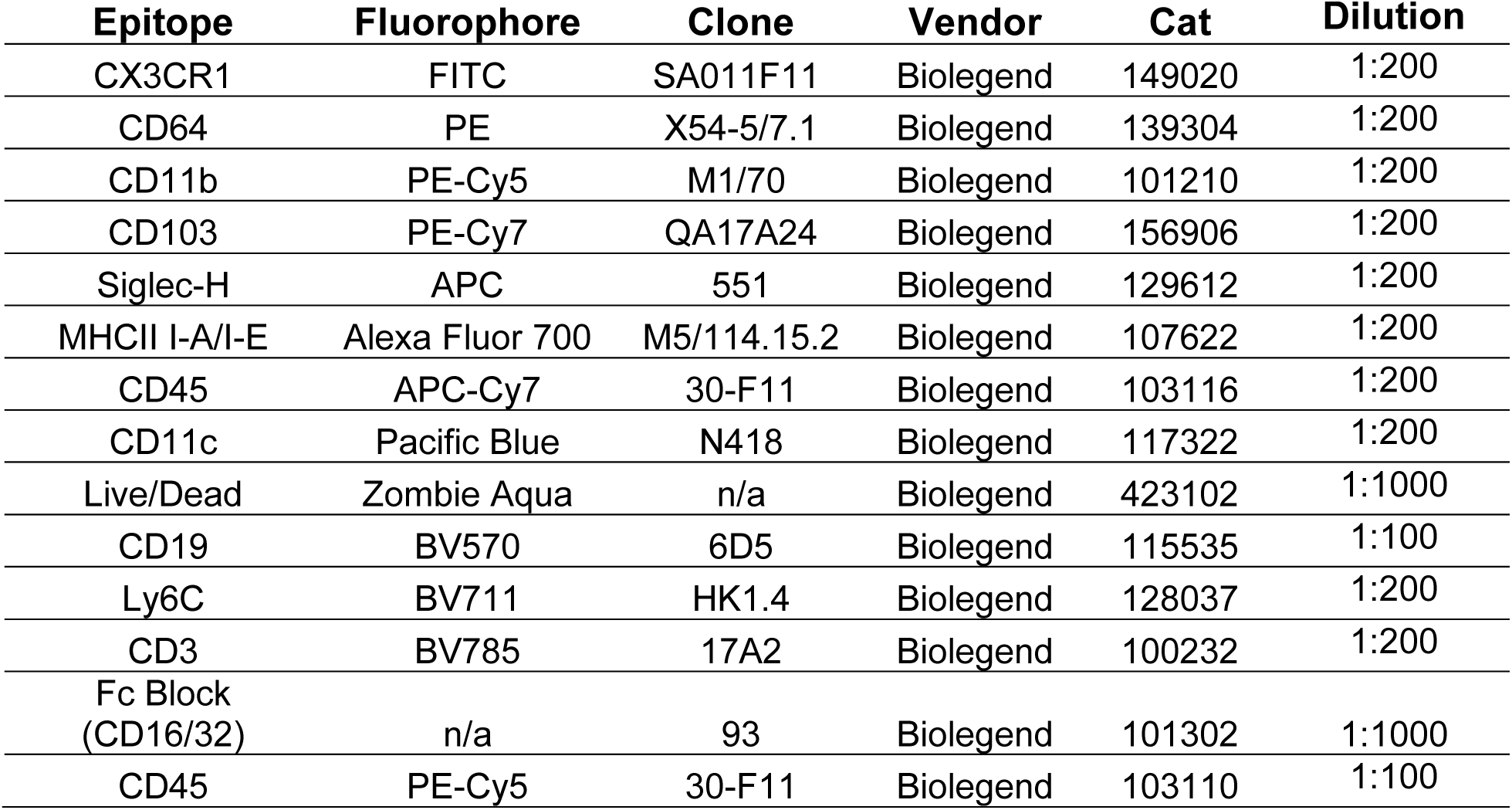
Antibody List.

Rhythmic analysis was performed using CircaCompare v0.2.0 to establish a best fit cosinor curve and provide analytical values for comparisons^29^.

### Fluorescence-Activated Cell Sorting (FACS)

Immune cells were washed, stained with Zombie Aqua viability stain, αCD16/32, and an αCD45 (PE-Cy5) antibody. Compensation and gating were performed with a mix of beads and unstained cells. The cells were then resuspended in FACS buffer and sorted 5 × 10⁶ live CD45+ immune cells with a BD FACS Aria for each time point, followed by downstream applications in the scRNA-seq protocol.

### Single Cell Suspension Preparation

Live CD45⁺ cells isolated via FACS were fixed and permeabilized in 4% formaldehyde buffer according to the 10x Genomics Chromium Fixed RNA Profiling protocol (Cat# PN-1000414). Cells were fixed for 22.5 hours, then spun down and resuspended in Quenching Buffer. Glycerol was added to reach a final concentration of 10%, and cells were stored at -80°C. FACS and fixation were performed in duplicate, and duplicate time points were pooled, resulting in 8 mice sequenced per time point for downstream analyses.

### Probe-Based Barcoding

Cells isolated from each of the four time points were hybridized with a 16-probe barcode set (PN-1000497, Probe Set v1.0.1). Cell viability and concentrations were assessed using the Countess II Cell Counter with AOPI viability dye (Fisher Scientific, NC2019981) to ensure equal pooling. Barcode and adapter ligations were performed by the McDermott Center Next Generation Sequencing Core at UT Southwestern Medical Center. Gel Bead-in Emulsion (GEM) generation was conducted using the Chromium Next GEM Chip and Chromium X instrument. Libraries were sequenced on an Illumina NovaSeq X Plus system using a 25B flow cell.

### Bioinformatic Analyses

Raw sequencing files were aligned to the UCSC mouse reference genome (mm10) using CellRanger. These raw sequence files are accessible through the NCBI BioProject Database under ID: PRJNA1405742. Features expressed in fewer than 100 cells were filtered out, along with cells containing fewer than 100 feature assignments or more than 10% mitochondrial transcripts. Since these single cell suspensions were subjected to FACS based on CD45 positivity, contaminating cells that were CD45 negative were bioinformatically filtered out as well as the few epithelial cells and endothelial cells that made up less than 0.3% and 0.05% of the entire dataset, respectively. After filtering, 815,073 high-quality live cells remained. All features were log-normalized and scaled. Due to dataset size, dimension reduction variables were computed using a representative subset of 50,000 cells. Integration was performed using the RPCA algorithm. The remaining ∼750,000 cells were projected onto the reduced-dimensional space using the ProjectIntegration() and ProjectData() functions from the Seurat package^30^. Data visualization was conducted using Seurat compatible with ggplot2^31^ and plotly^32^ R packages. Cells were annotated using marker genes generated via a negative binomial test in 10X LoupeR^33^.The MetaCycle^34^ R package was used to calculate oscillation scores and identify rhythmic features from the single-cell dataset. Trajectory analysis to uncover marker genes for differentiation was performed with Monocle3^35^. Cell communication through paired ligand-receptor analysis used CellChat R package^36^.

## Results

### Profiling small intestine immune cells using flow cytometry reveals oscillatory cell types, but with limited resolution

To profile small intestine immune cells across a day-night cycle, we harvested mouse lamina propria cells, including Payer’s patches, at several time points and evaluated them using flow cytometry and scRNA-seq experiments (**Fig.1A, Fig. S1A**). We used time coordination with Zeitgeber time (ZT) through a 12 hour light/12 hour dark cycle, where lights turn on at ZT0 and lights turn off at ZT12 over 2-3 weeks of acclimatization prior to cell harvest^37^.

**Figure 1.**
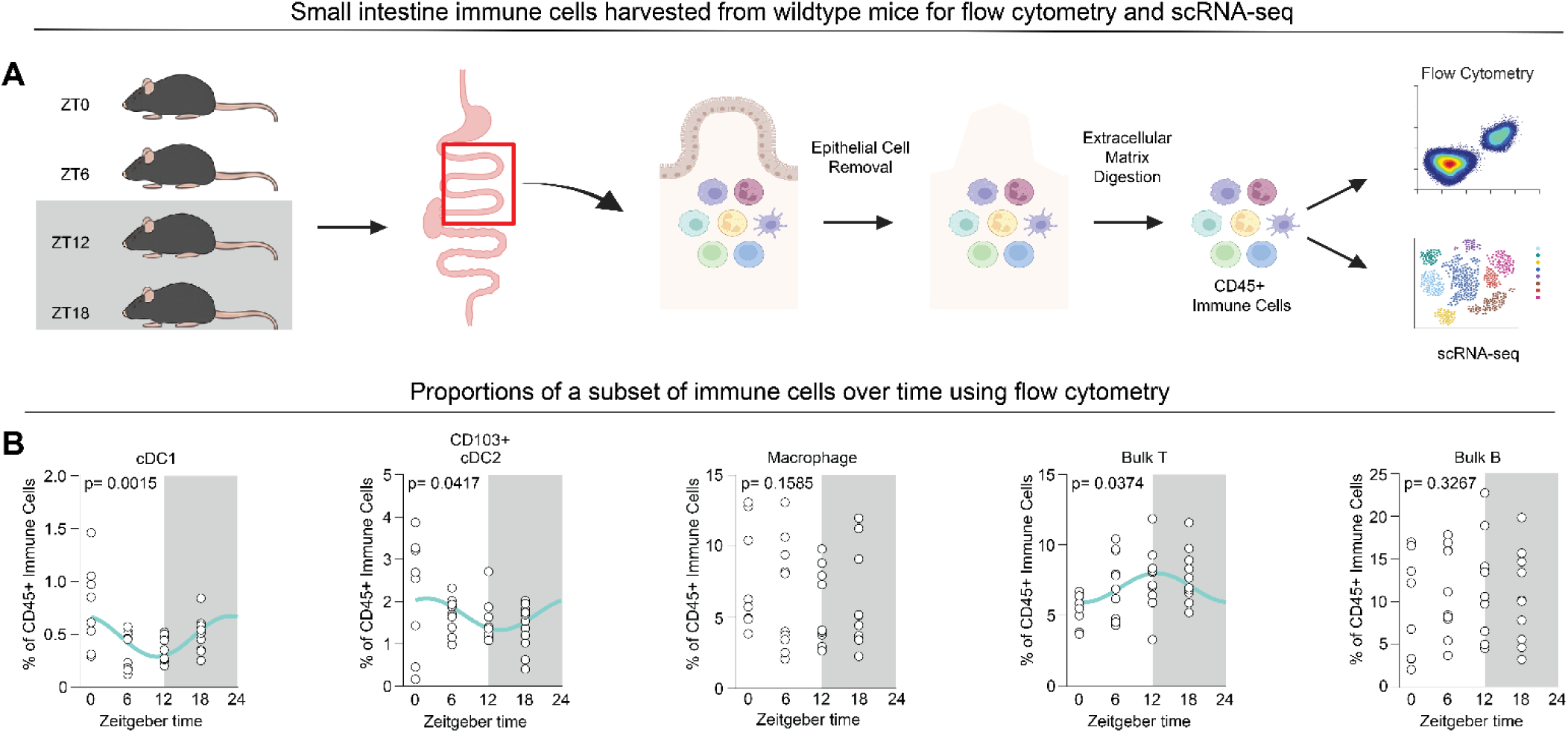
Experimental design for small intestine immune cell analysis over time and proportions of a subset of immune cells using flow cytometry. (A) Experimental design for capture of lamina propria immune cells and analysis using flow cytometry and scRNA-seq. Mice were acclimatized in light cabinets and at ZT0, 6, 12, or 18 small intestines were harvested, mechanically and enzymatically processed, and immune cells were isolated prior to flow cytometry or scRNA-seq. (B) Immune cell profiling over time using flow cytometry. A gating strategy (**Supp. Fig.1A**) using 12 antibodies quantified 10 cell types (**Supp. Fig.1B**), with absolute counts shown in **Fig.S1C**. Here, a subset of cell types are shown as the percent of CD45+ immune cells, with statistical analysis from CircaCompare. p-values are reported on the graph and the cyan line denotes the statistically significant (p=<0.05) best-fit nonlinear sinusoidal curve. n= 8-11 from two independent experiments.

We evaluated whether there were time-dependent differences in lamina propria immune cells using a flow cytometry gating strategy shown in **Fig.S1A**. We quantified absolute cell counts (**Fig.S1B**) and the percentage of live CD45+ immune cells (**Fig.1B**) for ten distinct cell types. Our 12-marker antibody panel focused primarily on myeloid cells, since myeloid cells rhythmically traffic via bone marrow egress^6^ and homing towards secondary lymphoid tissues^7,8^. To determine whether significant differences in 24-hour rhythmicity were present, we used CircaCompare, a rhythmicity detection program^29^. Subtypes of DCs and T cells showed differential proportional abundances across time (**Fig.1B**) but this was not the case for mature macrophages or B cells. Thus, immune cells in the small intestine show differences dependent on time. However, the relatively small number of protein-based cell markers used here limited resolution for broad immune cell annotation. Therefore, while flow cytometry is useful for focused panels targeting distinct cell types, it was insufficient to capture the vast breadth of immune cells in the small intestine for our purposes here.

### Single cell RNA sequencing provided the depth and resolution to examine the intestinal immune cell landscape over time

To gain deeper resolution of the entire breadth of immune populations, we performed scRNA-seq on lamina propria immune cells. Single-cell suspensions were harvested from the small intestine of eight mice at four distinct time points and we sorted 5 × 10⁶ live CD45⁺ cells. Following quality control filtering, 815,073 cells were retained for downstream analysis (see **Methods** and **Fig. 2A**). Cells were annotated using marker genes that we compiled based on 165 publications spanning 1981 to 2026 (**Supp. Table 1**). These marker genes revealed 33 distinct immune cell types, including 12 B cell subsets and 8 myeloid populations (**Supp. Table 1**, **Fig. 2B, Supp. Fig. 2A-B**).

**Figure 2.**
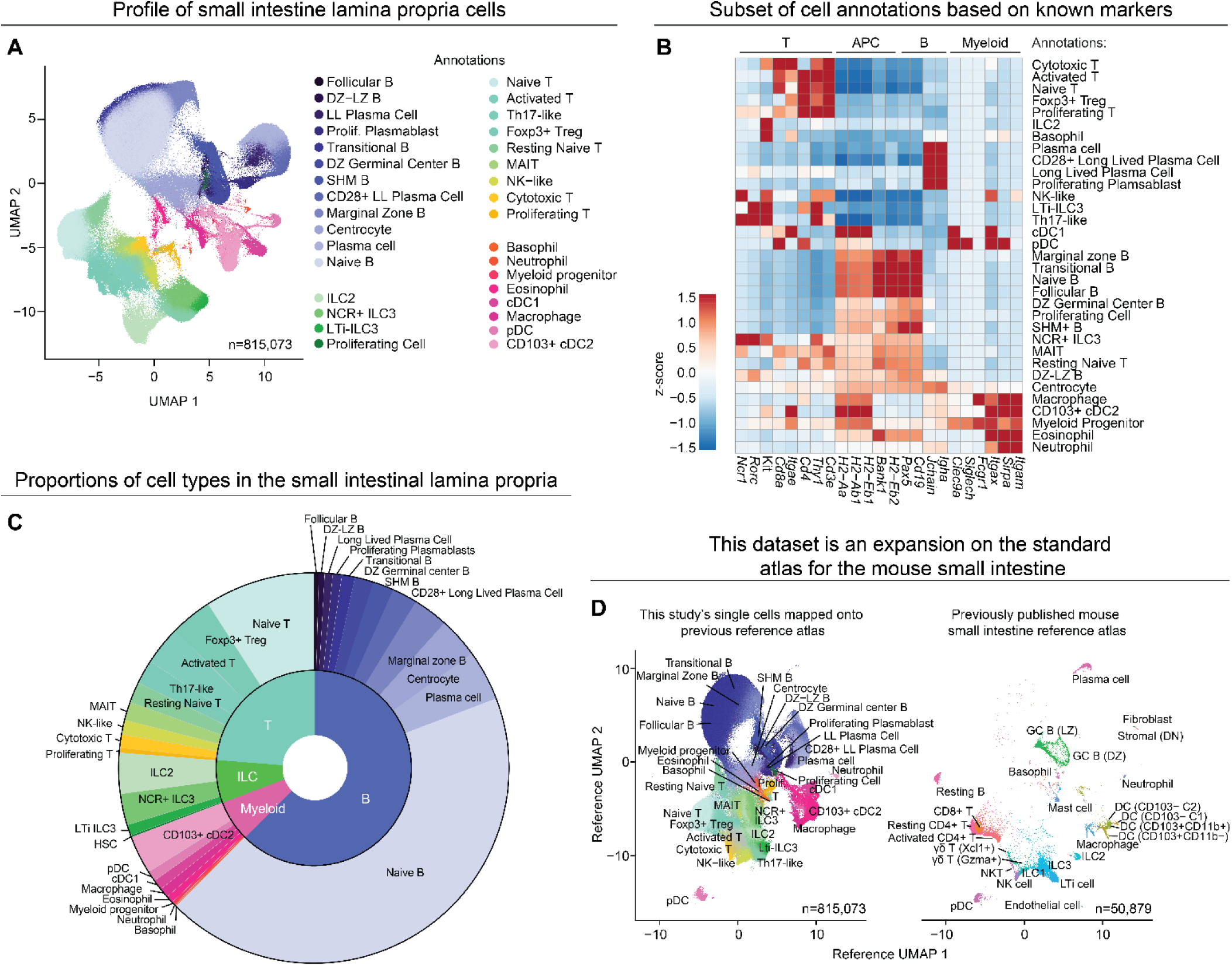
scRNA-seq of small intestine lamina propria immune cells establishes a broad atlas of over 30 cell types. (A) A UMAP plot of 815,073 immune cells that met quality control criteria from scRNA-seq, combining data from all four time points (ZT0, 6, 12, 18; n=815,073). cells were sequenced from two independent experiments with 4 mice for each time point. Abbreviations: Dark zone-light zone B cells (DZ-LZ B); Dark zone B cells (DZ B); Somatic hypermutation B cells (SHM B); Innate lymphoid cell (ILC); Natural cytotoxicity receptor (NCR); Long lived (LL); Plasma cell (PC); T helper (Th); Regulatory T cell (Treg); Mucosal-associated invariant T cell (MAIT); Natural killer (NK); Conventional dendritic cell (cDC); Plasmacytoid dendritic cell (pDC). (B) A z-score scaled heatmap for an abbreviated/condensed list of marker genes used to annotate the 33 clusters, separated by cell-type groups. Expanded heatmaps for hallmark genes are shown elsewhere for B cell markers (Fig. 5A), myeloid markers (**Supp.** Fig. 1A), and T and ILC markers (**Supp.** Fig. 1B). **Supp. Table 1** lists key marker genes and references for all annotations. (C) Sunburst chart showing the proportions of the annotated clusters within their respective cell-type groups out of all 815,073 cells. (D) A reference UMAP of previously published large-scale scRNAseq of the lamina propria (right)^40^. This dataset queried onto the reference UMAP of Xavier et al. with cell annotations labeled (right).

We observed a substantial proportion of naïve adaptive immune cells within the lamina propria at all time points. B cells were the most abundant cell type in the lamina propria and a majority of B cells expressed IgD and IgM, consistent with a naïve B cell phenotype (**Fig. 2C**, **Supp. Fig. 2C-D**). Another notably abundant population was naïve T cells, which express *Cd4* or *Cd8a* (**Fig. 2C, Supp. Fig. 2E**). Naïve B and T cells consisted of over half of the total small intestinal immune cells—a large but unsurprising amount considering the intestine is a key site for immune cell priming after exposure to microbiota and food antigens^38,39^.

We compared our dataset to another atlas of 58,067 mouse small intestinal lamina propria cells from a single time point^40^ (**Fig. 2D**). Our high yield dataset (815,073 cells) confirmed many canonical immune populations but also revealed rarer immune cell populations including subsets of transitional B cells, follicular B cells, and rare myeloid cells. Altogether, this dataset expands on previous profiling and creates comprehensive and kinetic single-cell immune maps of the intestinal lamina propria.

### Immune cell subpopulations of the lamina propria are dynamic and diurnally rhythmic

We next examined diurnal (24 hour) shifts in immune cell proportions and rhythmic gene expression. As a control, we first measured marker gene expression within each respective cell type to confirm that per-cell marker gene expression remained stable across time. This ensured that cell-type assignments were not influenced by time-of-day effects (**Supp. Fig. 3**). We then assessed how the sizes of these populations changed over the diurnal cycle. All major immune cell types — T cells, B cells, myeloid cells, and ILCs — displayed time-dependent fluctuations in abundance (**Fig. 3A**).

**Figure 3.**
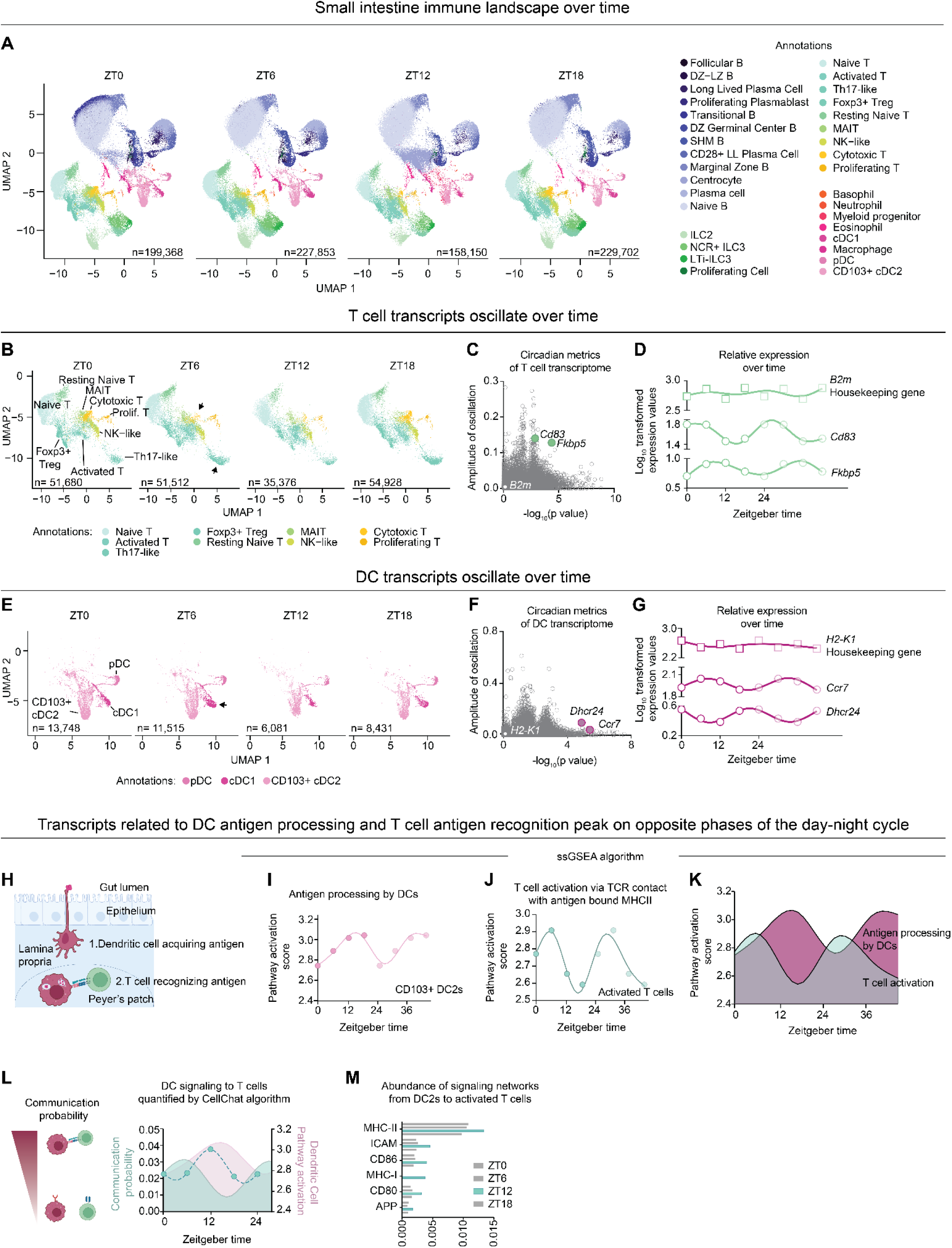
Analysis of immune cells over time reveals dynamic DC and T cell populations. (A) UMAPs of annotated cells separated by time of collection (ZT0, 6, 12, and 18). (B) Zoomed in UMAPs of T cells, separated by time of collection. Clusters that demonstrated significant alterations are identified with an arrowhead. (C) Scatter plot of the amplitude and -log(p-value) of the oscillation of normalized, scaled, and aggregated gene expression from T cells as measured by Metacycle. Select examples representing oscillatory (*Fkbp5, Cd83*) or non-oscillatory (*B2m*) genes are highlighted. (D) Individual plots of oscillatory (*Fkbp5, Cd160*) or non-oscillatory (*B2m*) gene expression within T cells over time. Time points are repeated for visualization purposes only. (E) Zoomed in UMAPs of dendritic cells, separated by time of collection. Clusters that demonstrated significant alterations are identified with an arrowhead. (F) Scatter plot of the amplitude and -log_10_(p-value) of oscillation of normalized, scaled, and aggregated gene expression from dendritic cells as measured by Metacycle. Select examples representing oscillatory (*Dhcr24, Ccr7*) or non-oscillatory (*Ly6e*) genes are highlighted. (G) Individual plots of oscillatory (*Dhcr24, Ccr7*) or non-oscillatory (*H2-K1*) gene expression in DCs across time. Time points are repeated for visualization purposes only. (H) Schematic of DC (pink) luminal antigen sampling and DC:T cell (green) contact. (I) Single sample gene set enrichment analysis of the antigen processing pathway (GO:0002604) of bulk CD103+ cDC2s over a 24 hour cycle. Data points repeated for visualization purposes only. (J) Single sample gene set enrichment analysis of the T cell activation pathway via antigen recognition (GO:0002291) of bulk activated T cells across a 24 cycle. Data points repeated for visualization purposes only. (K) Comparison of pathway scores for antigen processing by DCs and T cell activation across time. Data points repeated for visualization purposes only. (L) Schematic of DC (pink):T cell (green) communication states and communication probability generated from CellChat overlayed with Figure 3K data. (M) cDC2-activated T cell signaling network scores arranged by time, highlighting a peak at ZT12.

We focused on T cells and observed a significant fluctuation of Th17 and cytotoxic T subsets, most notably at ZT12 (**Fig. 3B**). We also calculated oscillation scores of T cells using MetaCycle, which assesses both a amplitude and statistical rhythmicity (**Fig. 3C, Supp. Table 2**). This analysis identified multiple oscillatory genes, including *Cd83*, a transmembrane glycoprotein upregulated in exhausted T cells^41^, and *Fkbp5*, a co-chaperone involved in glucocorticoid receptor-mediated stress responses and potential target for stress-induced circadian disruption^42,43^ (**Fig. 3D**). We also identified potential “housekeeping genes” that could serve as stable references for future temporal analyses. *B2m*, encoding beta-2-microglobulin, emerged as a strong candidate housekeeping gene, with consistently high expression across all time points and a lack of significant rhythmicity in T cells (**Fig. 3D**, **Supp. Table 2**).

Similarly, we examined changes in DCs over time, as prior studies have suggested their abundance fluctuates significantly throughout the day^7,8,44^. Among the DC populations, conventional dendritic cells type 1 (cDC1s) had the most pronounced diurnal variation in both abundance and UMAP-defined transcriptional states across the four time points (**Fig. 3E**). We further assessed their transcriptional profiles and identified a number of oscillatory genes, including two antiphasic, significantly oscillatory genes: *Dhcr24* and *Ccr7* (**Fig. 3F-G**). *Dhcr24* encodes 24-dehydrocholesterol reductase, which interacts with MAVS and STING in DCs, dampening antiviral immune responses^45^. *Ccr7*, a chemokine receptor for homing to secondary lymphoid tissues, also exhibits rhythmicity similar to DCs in skin draining lymphatics^8^. In addition, we identified candidate housekeeping genes for DCs based on consistently high expression and low oscillatory amplitude over time, including *H2-K1,* a Major Histocompatibility Complex II (MHC II) gene (**Fig. 3G** and **Supp. Table 3**).

Mice have temporally regulated immune responses to pathogens^1,2^, leading us to examine whether different cell types coordinate antigen recognition and response by time of day. Specifically, we used single-sample gene set enrichment analysis (ssGSEA) to determine whether the rhythmic transcriptional programs in antigen-presenting DCs were coordinated with T cell antigen responses (**Fig. 3H**). Our dataset suggested that antigen processing activity in CD103⁺ cDC2 cells peaks at ZT12, coinciding with the onset of activity and feeding in mice (**Fig. 3I)**. In contrast, transcripts related to T cell activation, including TCR signaling and response to MHC class II presentation, peaked 6–12 hours later (**Fig. 3J)**. This temporal offset indicates possible phased coordination: DCs are transcriptionally primed to capture and process antigen during the early active phase, while T cells transcribe their most robust activation gene responses in the subsequent rest phase (**Fig. 3K**). Supporting this model, paired ligand-receptor analyses demonstrated that communication probability between DCs and T cells also peaked at ZT12 (**Fig. 3L-M**). This lag may serve as a regulatory mechanism to optimize antigen presentation or may allow time for dendritic cell migration to a Peyer’s patch before full T cell engagement. Overall, our data suggest the temporal coordination and temporal delay of DC antigen processing and T cell activation, with cell interactions occurring as mice wake and interact with the environment.

### Small intestine immune cells have variable circadian gene expression

To complement our analysis of diurnal changes in immune cell abundance, we next assessed the magnitude and rhythmicity of the circadian clock genes themselves. We plotted the expression of circadian clock genes (KEGG pathway, mmu04710) and their amplitude of oscillation (**Fig. 4A**). Circadian genes drive transcription-translation feedback loops which establish the oscillatory gene expression (**Fig. 4B**). Circadian gene expression differed by cell cluster, with high expression in Th17s and ILCs (**Fig. 4A**, thick vertical grey lines), and low expression in terminally differentiated B cells and plasma cells (**Fig. 4A**, thin vertical grey lines). High expression and oscillation of circadian genes in Th17s and ILCs was expected given that these genes are necessary for their differentiation^9–12,46^. However, we found that terminally differentiated B cells—plasmablasts and plasma cells—have minimal circadian gene expression.

**Figure 4.**
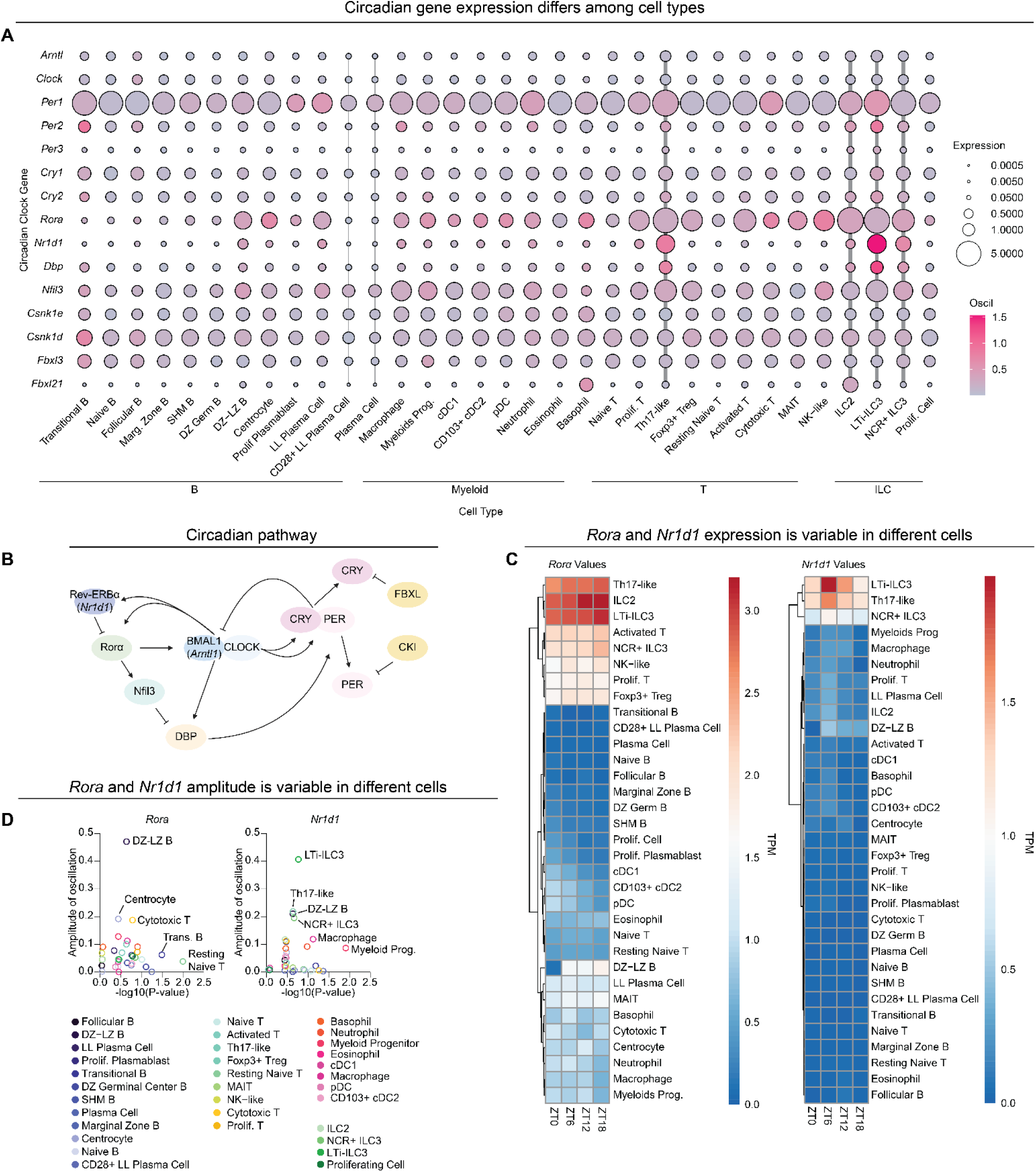
Circadian gene expression variability in small intestine immune cells. (A) Dot plot of circadian genes from KEGG Pathway (mmu04710) by cluster. Dot size represents gene TPM expression and color scale represents the amplitude of oscillation score generated from MetaCycle. Vertical gray lines denote clusters with high (thick) or low (thin) oscillation. (B) Schematic of the circadian transcription-translation feedback loop. For proteins with distinct gene names, gene names are provided below in italics. (C) Heatmaps of circadian genes *Rora* and *Nr1d1* with clustering from dendrograms showing TPM values across time. (D) Amplitudes of *Rora* and *Nr1d1* by log10 transformed p-values generated from MetaCycle.

To gain more insight into how circadian genes may influence specific immune cell subsets, we evaluated expression of individual circadian genes. Expression of certain circadian genes like *Clock* did not vary or oscillate much across distinct cell types (**Fig. 4A**). However, *Rora* and *Nr1d1* expression varied between cell types. We calculated average normalized expression for *Rora* and *Nr1d1* at each time point over a 24-hour cycle **(Fig. 4C)** and we calculated oscillation scores using MetaCycle, assessing both amplitude and statistical rhythmicity (**Fig. 4D**). *Rora* and *Nr1d1* expression was high and oscillatory in Th17s and ILCs, but expression was minimal in several cell types, including several B cell types (**Fig. 4C/D**). Taken together, expression of circadian genes differs between cell types, and can differ over the course of immune cell developmental stages.

### Variable circadian and immunoglobulin gene expression across B cell differentiation in the small intestine

B cells undergo selection and maturation in bone marrow and can then traffic to secondary immune tissues such as the intestine for further maturation and differentiation. Within the small intestine, Peyer’s patches serve as secondary lymphoid tissues to further modify B cell antibody affinity and isotype^47^. After encountering antigen, naïve B cells proliferate in the germinal center of the lymphoid follicle, cycling between dark zones (DZ) and light zones (LZ), and undergo somatic hypermutation and class switch recombination of immunoglobulin heavy chain genes to produce different antibody isotypes (IgG_1-4_, IgE, or IgA)^47^. Failure to develop a higher affinity antibody yields cell death via apoptosis. After class switching, B cells rearrange immunoglobulin light chain genes and terminally differentiate into antibody-producing plasma cells or memory B cells. Antibodies minimally consist of a heavy chain and light chain (**Fig. 5A** inset schematic).

**Figure 5.**
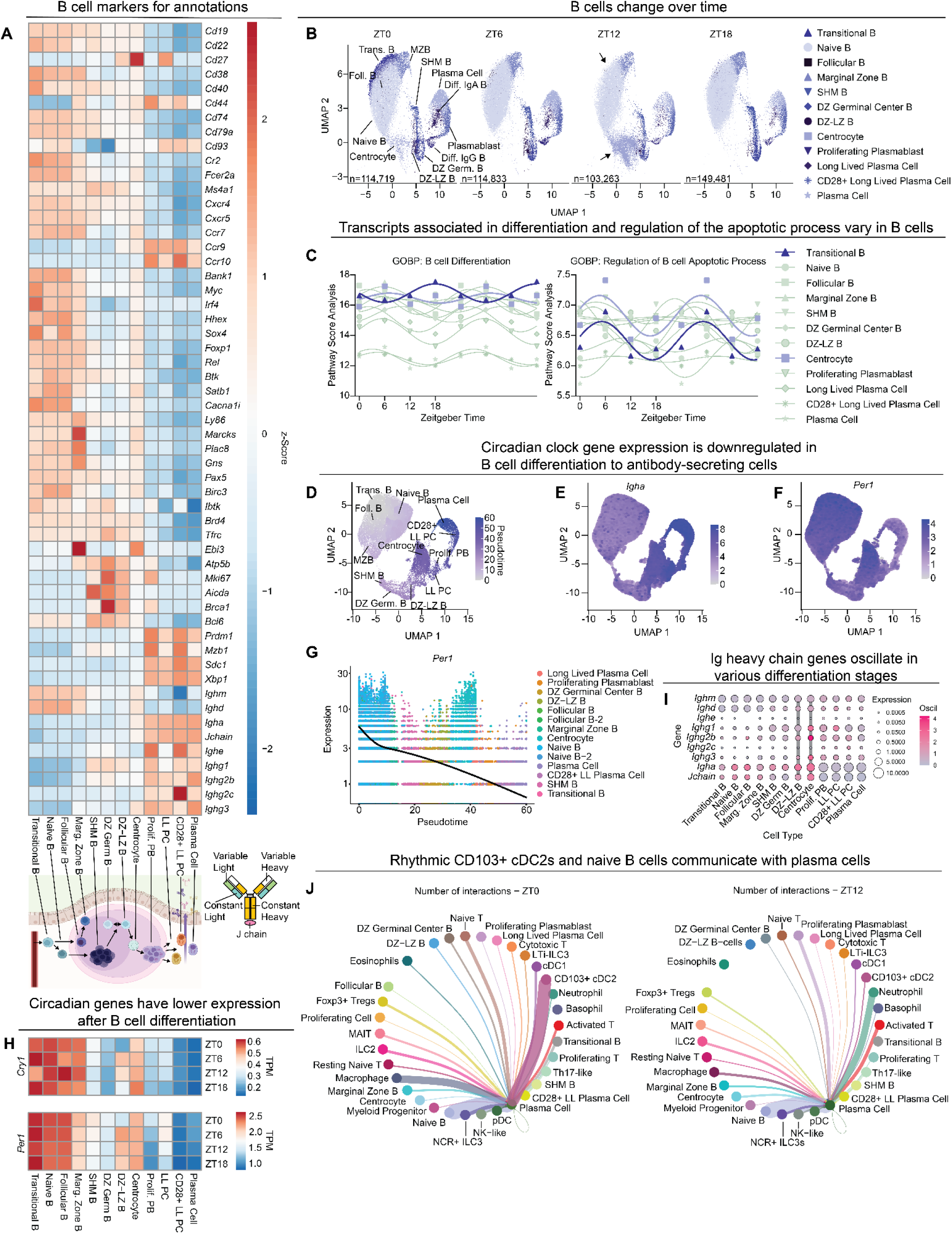
B cell clusters exhibit variability in gene expression and cell population dynamics dependent on differentiation stage. (A) Heatmap of hallmark genes used to annotate B cell clusters, arranged from early (left) to terminal (right) differentiation. Below are schematics of the differentiation process and structure of an antibody. (B) UMAPs of B-cell clusters (n=509,294) over time showing population differences. Arrows indicate populations with profound changes over time, centrocytes and transitional B cells. (C) ssGSEA analysis on Gene Ontology Biological Pathway analysis for B cell differentiation across time within B cell clusters, double plotted for visualization. (D) UMAP of trajectory analysis for B cell differentiation. Pseudotime begins with transitional B cells and ends with plasma cells. (E) UMAP of B cell trajectory analysis showing expression of *Igha*. (F) UMAP of B cell trajectory analysis showing expression of *Per1*. (G) Plot of *Per1* expression across B cell differentiation with the black line representing the average expression across pseudotime trajectory analysis. (H) Heatmaps of TPM values for circadian genes *Cry1* and *Per1*, arranged by differentiation stage and time. (I) Dot plot of immunoglobulin chain genes by B cell clusters. Dot size represents gene TPM expression; color scale represents the amplitude of oscillation score generated from MetaCycle. Vertical gray lines denote clusters with high oscillation. (J) Chord diagram showing plasma cell communication networks at ZT0 vs. ZT12. The line thickness denotes weight of prediction.

We annotated the B cell cluster and distinguished progressive stages of B differentiation via marker gene expression (**Fig. 5A, Supp. Table 1**). Annotations were informed by several genes, including immunoglobulin heavy chain genes from early stages in differentiation (*Ighm* and *Ighd),* hypermutation/class-switching and DNA repair genes (*Aicda* and *Brca1*), and a transcription factor required for secretion of immunoglobulins from differentiated antibody secreting cells (*Xbp1*)^48^. Additional genes informed annotations further, including factors involved in homing towards secondary lymphoid follicles with roles in early differentiation (*Ccr7, Cxcr4,* and *Cxcr5*) ^49^ and genes encoding proteins involved in retention within the lamina propria after terminal differentiation (*Ccr9 and Ccr10*)^50,51^.

To understand fluctuations among B cells, we examined shifts in cell proportions across time (**Fig. 5B**). Similar to other cell types, marker gene expression used to identify these B cell lineages was constant over time (**Supp. Fig. 3C**), so changes in cell populations are unlikely due to changes in marker gene expression. Two clusters had particularly profound changes over time—centrocytes and transitional B cells.

Transitional B cells peaked at ZT0 but were largely absent at all other time points (**Fig. 5B**, top black arrow) suggesting these immature B cells that originate from the bone marrow are highly dynamic in the small intestine. Centrocytes peaked at ZT12 but were essentially absent six hours before or after ZT12 (**Fig. 5B**, bottom black arrow).

Centrocytes differentiate from dark zone-light zone B cells (DZ-LZ B), raising the possibility that their proportional changes in the intestine vary diurnally. To correlate changes in centrocyte or transitional B cell populations to functional pathways, we performed ssGSEA analysis. We found that centrocytes and transitional B cells had high and sometimes oscillatory pathway scores for chemotaxis **(Supp. Fig. A)**, B cell proliferation **(Supp. Fig. 4A)**, B cell differentiation **(Fig. 5C)**, and regulation of B cell apoptotic processes **(Fig. 5C)**. These results suggest that populations of certain B cells shift dramatically by time of day, perhaps through cell migration to or from the small intestine, differentiation within the lamina propria, or cell death.

Since our dataset captured B cell states along their maturation trajectory, we expanded our analysis to examine transcriptional changes that span the full continuum of differentiation. Using a pseudotime-based algorithm (Monocle3), which computationally orders cells along an inferred differentiation trajectory, each B cell was positioned along an inferred developmental progression, which began at transitional B cells and ended with plasma cells as the terminally differentiated cell type (**Fig. 5D**).

This analysis identified genes whose expression patterns predict a cell’s stage within the B cell differentiation pathway (**Fig. 5E, Supp. Table 4**). Immunoglobulin genes and circadian genes were significant predictors of B cell differentiation progress. Expression of *Igha*, which encodes the IgA heavy chain, increased throughout differentiation, as expected (**Fig. 5F)**. *Per1*, a representative circadian gene, decreased in expression throughout differentiation toward plasma cells (**Fig. 5G**), confirming data in **Figure 4A**.

Given reduced expression of circadian clock genes in terminally differentiated B cells such as plasma cells (**Fig. 4A**, **Fig. 5F**), we further examined expression of circadian genes *Cry1* and *Per1*, the initial repressors in the feedback loop (**Fig. 4B**), in the B cell subsets. Expression of *Cry1* and *Per1* was high in B cells early in the differentiation process, including naïve, transitional, and marginal zone B cells (**Fig. 5H**). However, their expression was low late in the differentiation process, including plasmablasts and plasma cells. Centrocytes and DZ-LZ B cells within germinal centers of Peyer’s patches represent unique intermediate stages; they are transitioning from somatic hypermutation of immunoglobulin genes or class-switch recombination to affinity-based selection. Expression of *Cry1* and *Per1* in DZ-LZ B cells was particularly variable **(Fig. 5H)**. Similarly, although there is an overall reduction of *Per1* expression across B cell differentiation, there was a spike of expression around pseudotime arbitrary value of 40 (**Fig. 5G)**, corresponding with dark zone-light zone B cells (DZ-LZ) and centrocytes. These results suggest that oscillatory gene expression in centrocytes and DZ-LZ B cells may add additional variability at a stage where diversification is important.

Because antibody production is a major function of B cells, we examined immunoglobulin gene expression over time. B cells undergo class switching to produce different antibody isotypes during differentiation and IgA is secreted rhythmically into the intestine^15^, so we anticipated changes in immunoglobulin gene expression over the course of differentiation and possibly by time of day. Immunoglobulin heavy chains (IgD, IgM, IgG_1-4_, IgE, or IgA) dictate the antibody class, effector functions, and contribute to antigen recognition (**Fig. 5A** inset schematic). Of the B cell subsets, centrocytes and DZ-LZ B cells displayed low but oscillatory expression of immunoglobulin heavy chain genes (**Fig. 5I**, thick vertical gray lines). Conversely, plasma cells displayed high but non-oscillatory expression of immunoglobulin heavy chain genes, particularly *Igha* that encodes the IgA heavy chain (**Fig. 5I**).

Curiously, plasma cells have low circadian gene expression and non-oscillatory expression of immunoglobulin genes in spite of past work showing oscillatory fecal IgA levels^15^: How are plasma cells producing oscillatory IgA if they are “off the clock”? One possibility is that plasma cells receive cues from another cell type that is “on the clock”. For example, although *Igha* transcription is not rhythmic, it is possible that rhythmic inputs from other cells may alter post-transcriptional processes such as translation or secretion, culminating in rhythmic production of IgA. To examine the potential for cell communication, we used paired ligand-receptor analysis to display potential plasma cell interactions with other cell types over time. At ZT0, the highest cell-cell communication potential for plasma cells included CD103+ cDC2 cells and naïve B cells (**Fig. 5J**, left, **Supp. Fig. 4B**), both of which are abundant at ZT0 (**Fig. 1B, 3E, 5B**) and have moderate to high expression of circadian genes (**Fig. 4A, 5E-G**). Previous work has shown CD103+ cDC2s are needed for IgA and IgG generation of antibody secreting plasma cells^52^. Interestingly, at ZT12, communication potential between plasma cells and CD103+ cDC2s declined dramatically, with highest plasma cell communication scores transitioning to neutrophils and naïve B cells (**Fig. 5J**, right). Neutrophils localize near plasma cells and support their survival^53^. These data suggest that if oscillatory production of IgA relies on extrinsic cues this may be due to cell-to-cell communication between conventional dendritic cells and plasma cells. Overall, our data reveal new insights into oscillatory and non-oscillatory processes in B cells.

## Discussion

The connection between the circadian clock and immune system has steadily grown in recent years and our study provides an expansive view of immune cells and gene expression in the mouse small intestine. This work provides an atlas of over 800,000 immune cells in the small intestine, detailed markers for immune cell annotation, oscillatory genes, non-oscillatory housekeeping gene candidates, and highlights changes in cell populations by time of day. Importantly, our profiling revealed new aspects of immune biology, including evidence for possible phase-shifted antigen processing by DCs versus antigen recognition by T cells. Additionally, we highlight cell types with the highest transcriptional expression of circadian genes, such as Th17 and ILCs, which are likely under strong circadian control. Our data also indicate profound changes in certain B cell populations by time of day, including centrocytes and transitional B cells. Finally, our data suggest rhythmic gene expression in early B cell differentiation, while terminally differentiated antibody-producing plasma cells have diminished circadian clock gene expression but nonetheless may be influenced by communication with other oscillatory cells.

Naïve T and B cells were the most abundant immune cells in the mouse small intestine, which aligns with immune cell development shaped by exposure to antigens at mucosal sites. Gut-associated lymphoid tissue (GALT), which consists of Peyer’s patches, isolated lymphoid follicles, and the lamia propria, is a hub for antigen processing and presentation due to its proximity to metabolites and microbiota^39,54^. Immune cells “educated” in the intestine can traffic to other body sites for immune and behavioral regulation^55–57^.

Our data suggested that DC-T cell interactions are influenced by time of day. DCs can simultaneously engage with multiple T cells and interact with over 500 T cells in an hour^58^. We found that transcripts related to antigen processing in DCs peaked before those related to T cell activation, suggesting a sequentially coordinated process. In skin, Ince et al. showed similar a sequential order of DCs homing toward skin lymphatics before T cells^44^. Additionally, various components of the T cell activation pathway are rhythmic in response to OVA vaccination challenge^28^. Thus, temporally coordinated DC-T cell processes may be broad throughout various tissues.

We found that lamina propria cell abundance varies by time of day, particularly for certain cell types. For example, transitional B cells are abundant at ZT0 and are largely absent at other times, and centrocytes are abundant at ZT12 but are largely absent six hours before or after. (**Fig. 5B**). These profound apparent changes in cell populations could reflect movement of cells in or out of the small intestine, cell death, or differentiation into a different cell type. Variable expression of chemokine receptors (**Fig. 5A**) and high but oscillatory pathway scores for chemotaxis **(Supp. Fig. 4A)** suggests that cell trafficking may contribute to differential abundance for at least some cell types. Similarly, pathway scores for regulation of apoptosis **(Fig. 5C)** and B cell proliferation **(Supp. Fig. 4A)** were oscillatory, indicating that cell death or proliferation/differentiation may influence B cell populations at different times. We observe many time-specific differences occurring at ZT12, coinciding with onset of the dark phase when nocturnal mice become active and begin to feed and interact with the environment. ZT12 may serve as an inflection point, with mice receiving cues from feeding, the environment, and microbiota changes^13,15^. It has been proposed that diurnal separation of immune development/differentiation pathways from immune effector pathways may aid overall homeostasis^59,60^.

Our data suggest distinct regulation of gene expression in B cells at early differentiation stages versus mature antibody-producing plasma cells. Naïve B cells have high expression of circadian clock genes which is reduced in differentiated plasma cells (**Fig. 5**). Plasma cell gene expression is minimally oscillatory, including the IgA heavy chain gene, *Igha*. However, Penny et al. found oscillatory production of fecal IgA protein that was maintained in mice with B cell-specific deletion of the clock transcription factor BMAL1^15^. Instead of regulation by circadian clock genes, IgA levels were dependent upon cues from microbiota, feeding, and metabolites^15^. Our data support and extend these ideas, suggesting that plasma cell communication with other cell types (**Fig. 5J**, **Supp. Fig.4B**) may contribute to rhythmic production of IgA. The primary function of plasma cells is to produce up to thousands of antibodies per second^61,62^; therefore, perhaps plasma cells deprioritize clock control in favor of mass antibody production and instead rely on other cells for regulatory cues that influence protein production and/or secretion.

Our study generates an immune cell atlas of the mouse small intestine over time and provides new insights into immune biology and regulation. Ultimately, future exploration of these cells may reveal insight into rarer populations masked by broad categorizations and serve as a guide into intestinal circadian immunobiology.

## Acknowledgements

We thank Alex Crofts, J. David Farrar, Anne Satterthwaite, and John Schoggins for helpful suggestions on the manuscript. We thank Alyssa Guzman and Angie Mobley of the UTSW Flow Cytometry Core for their assistance in FACS. Lastly, we thank Vanessa Maddox and Jade Allen of the McDermott Sequencing Core, and Amir Segev (10X Genomics) for their guidance and assistance with scRNA sequencing.

---

**Supplemental Figure 1.**
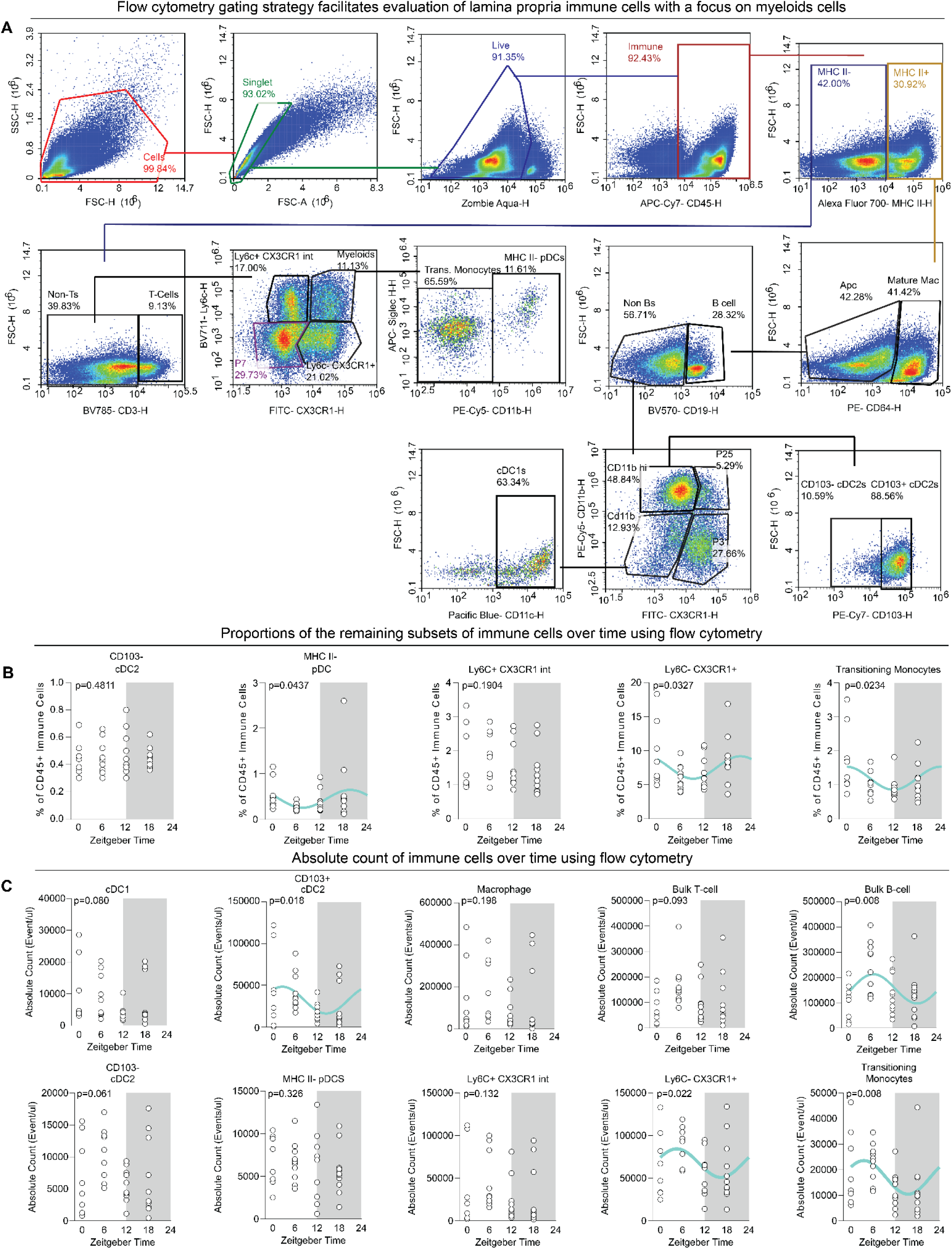
Flow cytometry gating strategy and absolute cell counts for lamina propria immune cells, related to Figure 1. (A) Representative flow cytometry gating strategy, with a focus on myeloid cells. Immune cells were gated as live CD45^+^ and then split into MHCII^+^ or MHCII^-^gates. MHCII^+^ cells were then gated by CD64 (CD64^+^ for Macrophages and CD64^-^ for remaining APCs). B cells were CD45^+^ MHCII^+^ CD19^+^ and non-B cells were further differentiated by Ly6C and CX3CR1 to distinguish between DCs subtypes. Bulk T cells were designated as CD45^+^ MHCII^-^ CD3^+^ with non- T cells subjected to further gating for MHCII^-^ myeloid cells. Ly6C^+^ CX3CR1^int^ were circulating monocytes and Ly6C^-^ CX3CR1^+^ were resident monocytes. Ly6C^+^ and CX3CR1^+^ myeloid cells were further gated to distinguish MHCII^-^ Ly6C^+^ CX3CR1^+^ CD11b^+^ Siglec H^+^ pDCs or MHCII^-^ Ly6C^+^ CX3CR1^+^ CD11b^+^ transitioning monocytes. (B) Other subsets of immune cells extended from **Figure 1B** with statistical analysis from CircaCompare. n= 8-11 from two independent experiments and p values are reported on the graph and the cyan line denotes the statistically significant (p=<0.05) best fit nonlinear sinusoidal curve. (C) Absolute count of lamina propria immune cells. n= 8-11 from two independent experiments with statistical analysis from CircaCompare. P values are reported on the graph and the cyan line denotes the statistically significant (p=<0.05) best fit nonlinear sinusoidal curve.

**Supplemental Figure 2.**
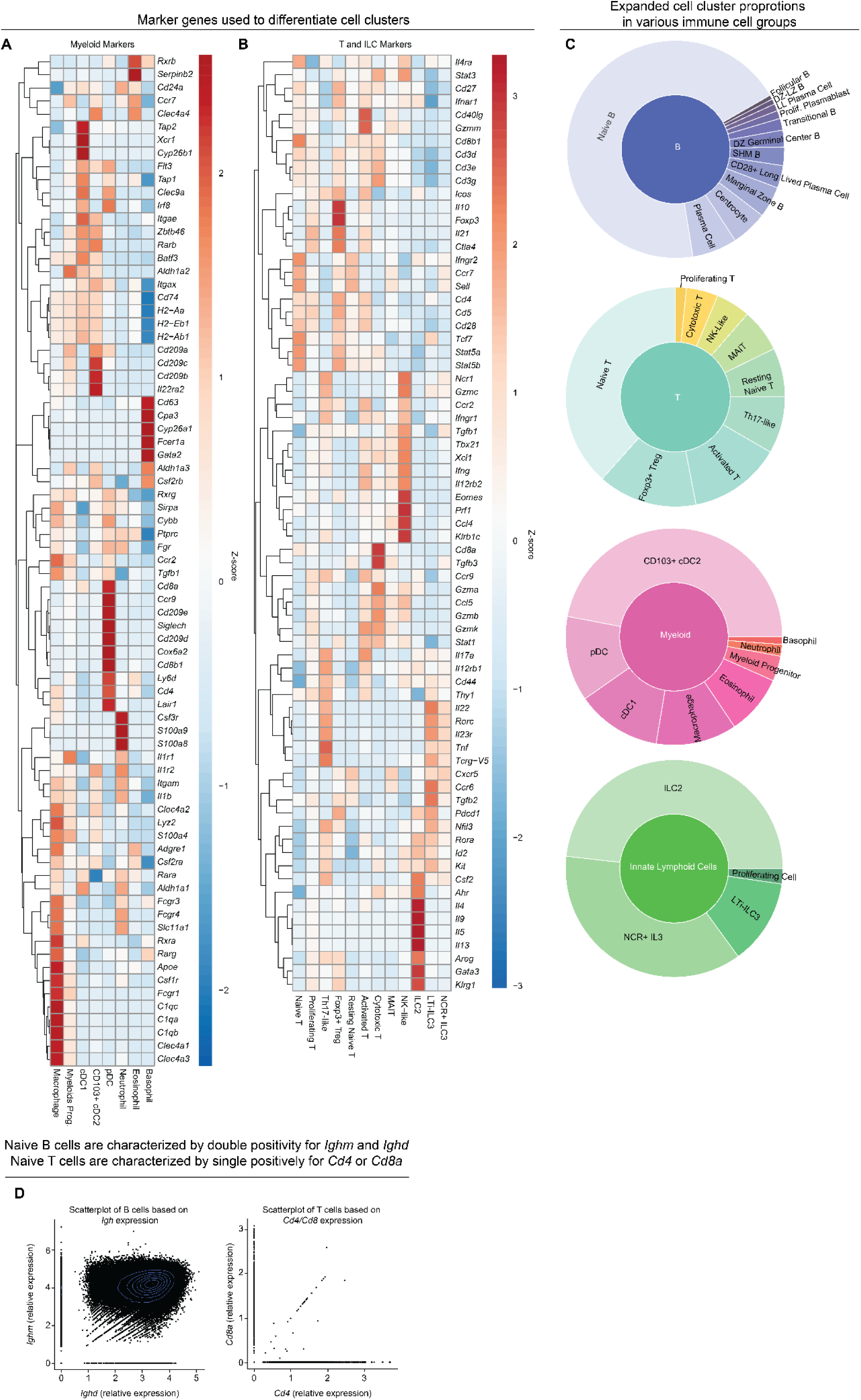
Extended markers used to differentiate clusters and expanded proportions of cell groups, related to Figure 2. (A) Heatmap of expanded list of makers with normalized expression used in annotating the myeloid clusters. Dendrograms show clustering arrangements. (B) Heatmap of expanded list of markers with normalized expression used in annotating the T cell and ILC clusters. Dendrograms show clustering arrangements. (C) Sunburst graph of expanded cell cluster proportions by cell type groups generated with Plot.ly R package. (D) Scatterplot graph evaluating double positive B and T cells by classifying genes

**Supplemental Figure 3.**
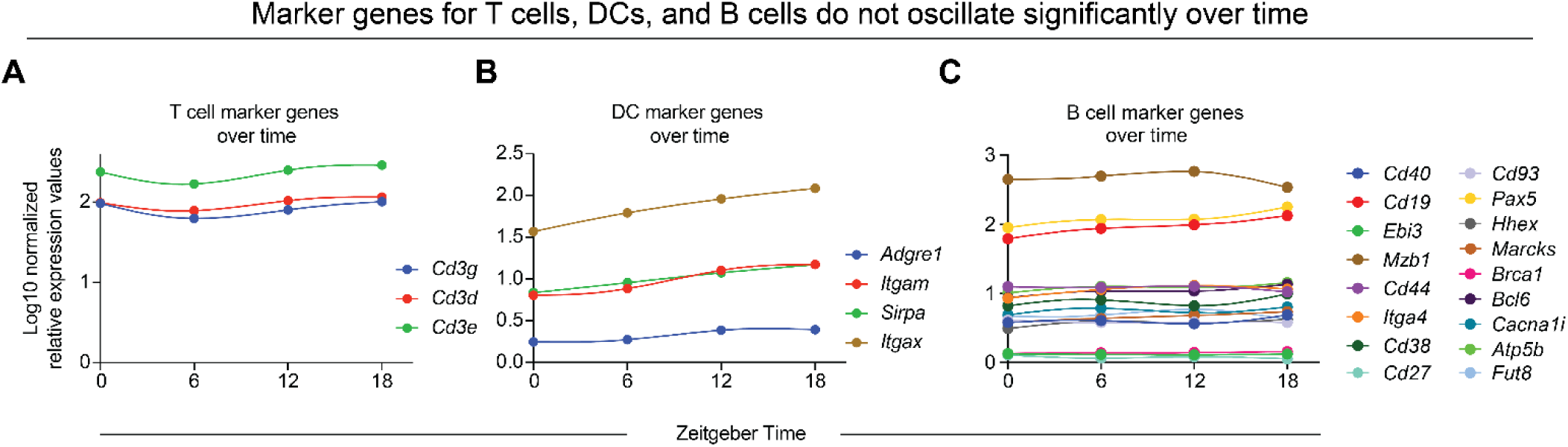
Marker genes from T cells, DCs, and B cells show minimal oscillation throughout the day-night cycle, related to Figure 3 and Figure 5. (A) Line graph of T cell marker genes for CD3 show minimal oscillation across time in Log10 normalized expression values. (B) Line graph of DC marker genes for various integrins, an adhesion molecule, and an inhibitory receptor show minimal oscillation across time in Log10 normalized expression values. (C) Line graph of B cell marker genes for integrins, surface molecules, transcription factors, and genes for B cell functionality show minimal oscillation across time in Log10 normalized expression values.

**Supplemental Figure 4.**
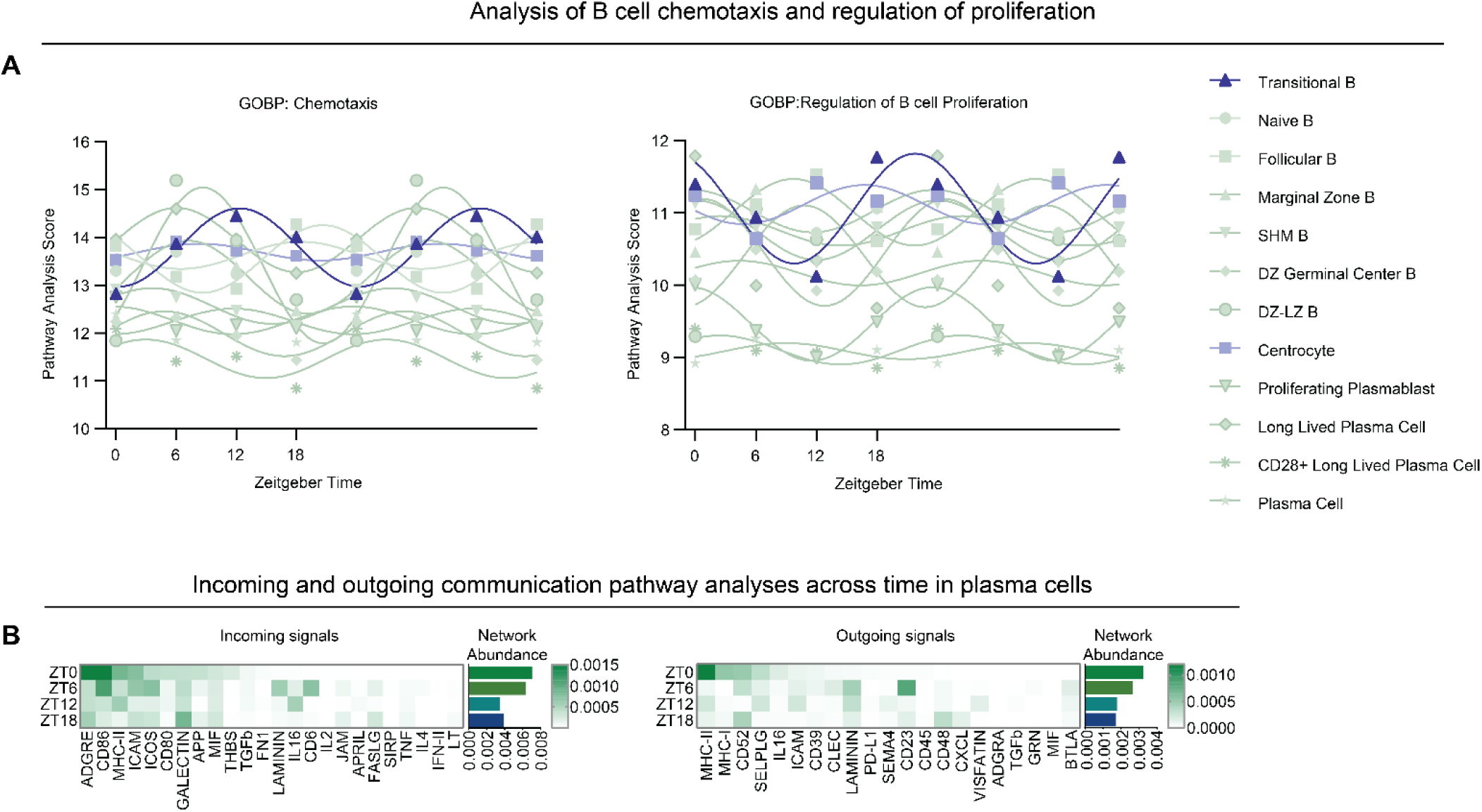
Extended Analysis of B cell clusters using Gene Ontology Biological Processes and breakdown of communication pathways from plasma cells, related to Figure 5. (A) ssGSEA analysis on Gene Ontology Biological Pathway for chemotaxis and regulation of B cell proliferation across time within B cell clusters, double plotted for visualization. (B) CellChat algorithm analysis of incoming and outgoing signals of paired ligand-receptor interactions of plasma cells across time. Network abundance highlights the sum of network interactions at the given time.

**Supplemental Table 1.**
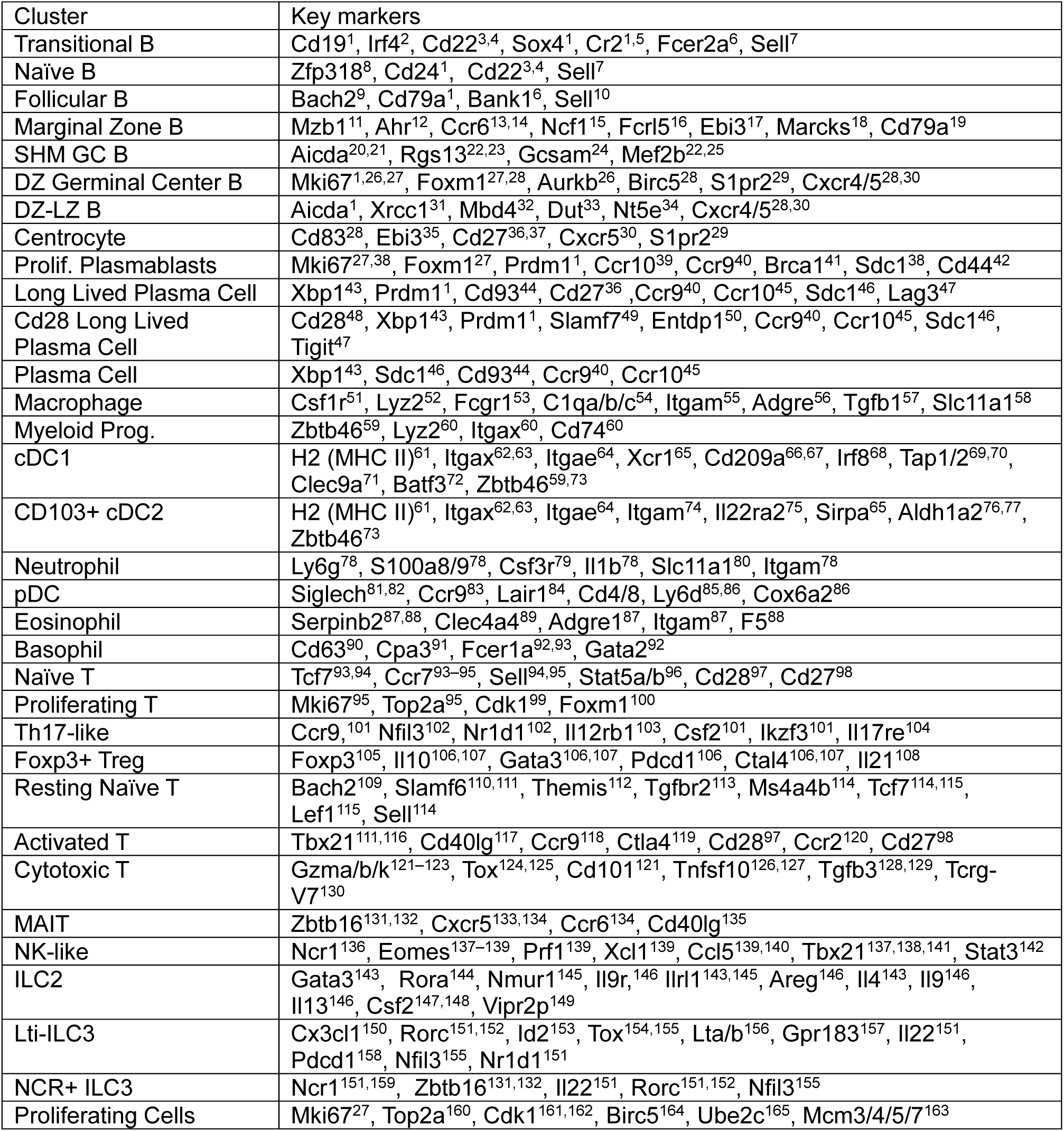
Key markers and references for cell cluster annotations.

